# Inflammation in schizophrenia: Peripheral interleukin-6-related disease-specific functional activity abnormalities

**DOI:** 10.64898/2026.02.06.704308

**Authors:** Yun-Shuang Fan, Jinxing Chen, Liju Liu, Cuiling Zhang, Jing Guo, Huafu Chen, Mi Yang

**Affiliations:** The Clinical Hospital of Chengdu Brain Science Institute, School of Life Science and Technology, University of Electronic Science and Technology of China, Chengdu, China; MOE Key Lab for Neuroinformation, Brain-Computer Interface & Brain-Inspired Intelligence Key Laboratory of Sichuan Province, University of Electronic Science and Technology of China, Chengdu, China

**Keywords:** Schizophrenia, Interleukin-6, Multimodal neuroimaging, Imaging transcriptomics, Immunity

## Abstract

**Background:** Peripheral inflammation is implicated in the pathophysiology of schizophrenia, but how inflammatory signals map onto the large-scale brain organization remains incompletely understood.

**Methods:** We applied a supervised multimodal fusion approach guided by interleukin-6 (IL-6) to gray matter volume (GMV) and resting-state regional homogeneity (ReHo) from a population-based discovery cohort in the UK Biobank. Brain components related to IL-6 were identified and then projected onto an independent schizophrenia cohort to examine their relevance to the disease. Imaging-transcriptomic analyses using the Allen Human Brain Atlas characterize the molecular substrates underlying the disease-relevant pattern.

**Results:** Two ReHo components were significantly associated with plasma IL-6, while no GMV components showed robust IL-6 correlations. One of the components (ReHo IC4) exhibited a conserved functional pattern characterized by enhanced visual synchrony and reduced synchrony in the medial prefrontal cortex. This pattern remained unchanged in both the healthy controls and patients. In contrast, another component (ReHo IC8) showed increased synchrony in the default mode network and reduced synchrony in sensorimotor networks, and its loadings were significantly elevated in patients with schizophrenia. Imaging-transcriptomic analysis revealed the molecular architecture of this disease-amplified pattern. The default mode region was enriched in synaptic signaling pathways, while the sensorimotor region was linked to mitochondrial bioenergetic processes; both patterns significantly enriched with gene sets related to schizophrenia.

**Conclusions:** This study identified an IL-6-associated functional brain pattern that is amplified in schizophrenia, linking peripheral inflammation to disease-specific network dysregulation. The findings provide a systems-level framework for understanding how peripheral inflammation interacts with large-scale brain network activities in schizophrenia.

## Introduction

Schizophrenia (SZ) is a common, severe psychiatric disorder characterized by extensive abnormalities in brain structure and function. Its etiology involves multiple factors, including genetic and environmental factors, such as immunity (Müller, 2018; Tandon et al., 2024). The traditional view holds that the brain is shielded from peripheral inflammatory responses by the blood–brain barrier. Recently, accumulating evidence has shown that inflammatory signals can access and influence the brain through various pathways (Upthegrove & Khandaker, 2019). These inflammatory signals modulate the function of neurons and glial cells, leading to brain alterations related to pathological states (Goldsmith et al., 2023). Nevertheless, the mechanisms underlying the association between inflammatory mediators and brain abnormalities in SZ remains poorly understood.

Interleukin (IL)-6, as a key mediator of immune and inflammatory responses (Rose-John et al., 2006), may serve as a biomarker of inflammatory status and disease progression in SZ (Chase et al., 2016; Zhilyaeva et al., 2023). Specifically, IL-6 can influence the structure and function of neurons, glial cells, and synapses (Kummer et al., 2021; Liu et al., 2022; Mirabella et al., 2021; Wei et al., 2011). Higher IL-6 levels have been observed in the peripheral blood and cerebrospinal fluid of patients with SZ (Orlovska-Waast et al., 2019; Zhou et al., 2021). Furthermore, Mendelian randomization and related genetic instrumental variable analyses have confirmed that a potential causal relationship between elevated IL-6 levels and the risk of SZ (Perry et al., 2021). Neuroimaging studies have reported that increased IL-6 levels are associated with gray matter atrophy in the middle temporal gyrus and fusiform gyrus, cortical thinning in the superior frontal gyrus (Williams et al., 2022), and functional hypoconnectivity between the medial prefrontal cortex (mPFC) and sensory processing areas in SZ (Mlynek et al., 2025). Overall, these findings emphasize the crucial role of IL-6 in the pathophysiology of SZ.

SZ is associated with cross-modal disruptions in coordinated brain networks, which suggests that IL-6 should be included as a reference variable in cross-modal neuroimaging analyses. Patients with SZ show a joint structure–function impairment pattern involving visual cortical functional activation, anterior thalamic radiation white-matter integrity, and parietal gray matter volume (Sui et al., 2013). A supervised multimodal fusion approach called multi-site canonical correlation analysis with reference plus joint independent component analysis (MCCAR+jICA) was used to identify target-related cross-modal components (Qi et al., 2018). Using MCCAR+jICA, researchers have identified specific brain components associated with certain biomarkers or phenotypic scores. For example, an Alzheimer’s disease study employed this strategy and discovered a structure–function covariation pattern related to neuropsychiatric symptom, which could further predict longitudinal cognitive decline (Li et al., 2023). Moreover, based on this method, a multimodal frontotemporal network has been found to be associated with SZ polygenic risk scores (Qi et al., 2022). Therefore, MCCAR+jICA can be used as an effective method to identify IL-6-related multimodal neuropathological features in SZ.

Genes serve as the blueprint for shaping brain organization and are involved in the etiology of SZ (Hibar et al., 2015). The Allen Human Brain Atlas (AHBA) microarray dataset enables researchers to identify brain transcriptomic profiles relevant to brain structure and function. This transcriptomic dataset has been widely used to identify gene sets associated with neuroimaging abnormalities in various pathological states (Dear et al., 2024; Negi & Guda, 2017; Sunkin et al., 2012). For instance, a study on bipolar disorder found that expression levels of 27 specific genes were correlated with an abnormal regional homogeneity (ReHo) pattern, indicating the genetic basis of abnormal brain activity (Lv et al., 2025). Furthermore, the spatial expression pattern of SZ risk genes can predict the spatial alterations in gray matter volume (GMV) in SZ patients (Forest et al., 2017). By integrating neuroimaging data with gene transcriptomics, we can better understand the genetic architecture underlying IL-6-related macroscopic brain patterns of SZ.

In this study, we used the UK Biobank (UKB) dataset to investigate IL-6–associated multimodal brain pattern deviations in SZ, and to further reveal its transcriptomic underpinnings (Error! Reference source not found.). To select inflammation-related components, we applied the supervised MCCAR+jICA approach on multimodal brain patterns of a healthy population with IL-6 as the reference vector. We then projected the IL-6-related patterns onto an independent clinical cohort, and tested group deviations of component loadings in SZ relative to healthy controls (HC). We further performed imaging–transcriptomic partial least squares (PLS) analysis and functional enrichment to investigate the genetic and biological contexts. We hypothesized that specific IL-6–associated brain components would exhibit significant group differences between SZ and HC.

## Material and Methods

### Participants

#### Discovery cohort (Healthy population)

For the discovery cohort, we used the UKB dataset, a large population-based research resource comprising extensive phenotypic, biological, and neuroimaging data from individuals aged 40–69 years. The acquisition of data was conducted across 22 UK-based assessment centers from 2006 to 2010 throughout the United Kingdom. Ethical approval for the UK Biobank study was granted by the North West Multi-Center Research Ethics Committee (Approval number: 11/NW/0382), and all participants provided written informed consent (Sudlow et al., 2015). The current analysis was approved under project No. 569519. After selecting individuals with available plasma IL-6, structural T1-weighted MRI, and resting-state functional MRI (rs-fMRI) data, a total of 827 participants were included in the present study. Individuals were excluded if they had any ICD-10-coded neurological or psychiatric disorders, congenital brain abnormalities, or incomplete neuroimaging or IL-6 data. All included participants were free from major neurological or psychiatric conditions and passed the UK Biobank imaging quality control procedures.

#### Application cohort (SZ dataset)

For the application cohort, we utilized the Center for Biomedical Research Excellence dataset (COBRE, http://fcon_1000.projects.nitrc.org/indi/retro/cobre.html). The COBRE cohort includes structural and rs-fMRI data collected from 72 patients with SZ and 74 demographically matched HCs, with ages ranging from 18 to 65 years. All participants provided written informed consent in accordance with the institutional review board of the University of New Mexico.

### Plasma IL-6 measurement

Plasma IL-6 levels were obtained from the Olink Explore 1536 proteomics dataset in the UKB. Protein quantification was performed using the Proximity Extension Assay technology, and expression values were reported as normalized protein expression (NPX), a log2-scaled and internally normalized unit generated by Olink’s MyData Cloud platform. All NPX data underwent standardized plate and batch normalization according to Olink’s quality control procedures. To minimize the influence of potential outliers, extreme IL-6 NPX values were further excluded using an interquartile range–based approach. The resulting IL-6 values were used as the reference variable in the following multimodal fusion analysis.

### Multi-modal neuroimaging data acquisition

#### Discovery cohort

Imaging data were acquired using a standard Siemens Skyra 3T scanner (Siemens, Erlangen, Germany) with a standard 32-channel radio-frequency receiver head coil. High-resolution T1-weighted structural images were acquired using a 3D magnetization-prepared rapid gradient-echo (MPRAGE) sequence. The acquisition parameters were as follows: repetition time (TR) = 2000 ms; echo time (TE) = 2.01 ms; inversion time (TI) = 880 ms; flip angle = 8°; field of view (FOV) = 208 × 256 × 256 matrix; and isotropic voxel resolution of 1.0 × 1.0 × 1.0 mm^3^. Functional MRI data were acquired using a gradient-echo echo-planar imaging sequence with multiband acceleration. Key parameters included: multiband acceleration factor = 8; TR = 735 ms; TE = 39 ms; flip angle = 52°; field of view (FOV) = 88 × 88 × 64 matrix; and isotropic voxel resolution of 2.4 × 2.4 × 2.4 mm^3^. The scan duration was approximately 6 minutes, comprising 490 volumetric time points with anterior-to-posterior (A >> P) phase encoding.

#### Application cohort

Imaging data were acquired using a standard Siemens TrioTim 3T scanner (Siemens, Erlangen, Germany). High-resolution T1-weighted structural images were acquired using a 3D multi-echo magnetization-prepared rapid gradient-echo (MPRAGE) sequence. The acquisition parameters were as follows: TR = 2530 ms; TE = 1.64 ms; TI = 900 ms; flip angle = 7°; FOV = 256 × 256 matrix; and isotropic voxel resolution of 1.0 × 1.0 × 1.0 mm^3^. Functional MRI data were acquired using a gradient-echo echo-planar imaging sequence. Key parameters included: TR = 2000 ms; TE = 29 ms; flip angle = 75°; FOV = 64 × 64 matrix; and voxel resolution of 3.75 × 3.75 × 4.5 mm^3^. The functional scan comprised 149 volumetric time points.

### Data preprocessing

Structural T1-weighted images were preprocessed using the CAT12 toolbox (v12.9) (https://neuro-jena.github.io/cat) integrated within SPM12(www.fil.ion.ucl.ac.uk/spm). Preprocessing steps included bias-field correction and affine registration, followed by tissue segmentation into gray matter, white matter, and cerebrospinal fluid. Spatial normalization was performed using the DARTEL algorithm, creating normalized images resampled to 1.5 × 1.5 × 1.5 mm^3^. To preserve the absolute amount of tissue, the GM images were modulated using the Jacobian determinants (linear and non-linear) from the deformation fields. Finally, a 3-mm FWHM Gaussian kernel was applied for spatial smoothing.

Functional data preprocessing was carried out using DPABI (http://rfmri.org/DPABI). The initial 10 volumes were discarded. The remaining data underwent slice-timing correction and realignment. Participants exhibiting head motion exceeding 3 mm in translation or 3° in rotation were excluded. Images were then normalized to the Neurological Institute (MNI) space and resampled to 3 × 3 × 3 mm^3^ voxels. Nuisance covariates, including Friston-24 motion parameters, white matter, and cerebrospinal fluid signals, were regressed out. Linear detrending and bandpass filtering (0.01–0.1 Hz) were applied to reduce physiological noise and drift. ReHo was calculated voxel-wise using Kendall’s Coefficient of Concordance based on the time series of a central voxel and its 26 neighbors. To normalize variations across subjects, individual ReHo maps were divided by the global mean ReHo. These standardized maps were smoothed with a 4-mm FWHM kernel. Age and sex effects were regressed out from both GMV and ReHo data.

### Fusion multimodal data with reference

To investigate IL-6-related multimodal brain components, we used a fusion-with-reference model namely MCCAR + jICA (Qi et al., 2018). Here IL-6 was set as the reference to guide the joint decomposition of two MRI features (GMV and ReHo) based on supervised learning. The correlations of imaging components with IL-6 were maximized in the supervised fusion method, as in **Eq. 1**.

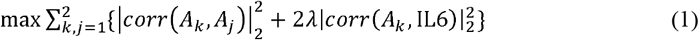

where *A*_*K*_ is the loading matrix for each modality, *corr*(*A*_*k*_, *A*_*j*_) is the column-wise correlation between *A*_*K*_ and *A*_*j*_, and *corr*(*A*_*k*_, IL6) is the column-wise correlation between *A*_*K*_ and IL-6. This supervised fusion method can extract joint multimodal components that are significantly associated with IL-6. This procedure yielded subject-specific loading coefficients for each modality, which were used for subsequent analyses.

### Component Selection and Characterization

Subject-specific loading coefficients for each independent component were obtained from the multimodal fusion model. In the discovery cohort, components whose loadings were significantly associated with plasma IL-6 after the false discovery rate (FDR) correction (*q* < 0.05) were designated as inflammation-related and were further used in subsequent analyses. For each significant component, spatial Z-maps were generated and thresholded at |*Z*| > 3 to visualize modality-specific contributions from GMV or ReHo. The spatial correspondence between modalities was examined to characterize the anatomical distribution of IL-6–associated multimodal patterns.

To study the relationship between inflammation-related brain patterns and brain abnormalities in SZ, the IL-6–associated components were linearly projected onto the independent SZ cohort. Group comparisons between individual loadings of SZ patients and HCs were performed using two-sample *t*-tests, with significance determined at FDR-corrected *p* < 0.05. Components showing significant group differences were considered both inflammation-linked and disease-relevant, and were subsequently used in the following analysis.

### Imaging-Transcriptome Association Analysis

Whole-brain microarray-based gene expression data were obtained from the AHBA (http://human.brain-map.org), comprising postmortem samples from six neurotypical adult donors (Hawrylycz et al., 2012). Gene expression values were preprocessed and averaged across donors following the standard *abagen* pipeline (https://abagen.readthedocs.io/en/stable) and aligned to the Schaefer-400 cortical parcellation(Arnatkevic□iūtė et al., 2019; Markello et al., 2021; Schaefer et al., 2018). The IL-6–related ReHo component map was resampled to the Schaefer-400 cortical atlas to generate a parcel-wise imaging vector. PLS regression was applied to identify the spatial correspondence between regional ReHo patterns and gene expression profiles, with gene expression as predictors and the parcel-wise ReHo as the response variable. The first latent component (PLS1) was retained for further analyses. Model significance was assessed using permutation tests (10,000 iterations), and the stability of gene weights was estimated using bootstrap resampling (1,000 iterations). Gene weights were converted to *Z*-scores, and genes with |Z| > 3 were selected for downstream functional enrichment. Gene Ontology (GO), Kyoto Encyclopedia of Genes and Genomes (KEGG), and disease enrichment analyses were performed using ToppGene (https://toppgene.cchmc.org), with statistical significance set at FDR-corrected *p* < 0.05.

## Results

### Participant characteristics

A final sample of 777 UK Biobank participants (59.79 ± 7.45 years; 55% female; demographic distributions shown in **Figure S1**) was included in our analysis after stringent quality control. An independent SZ cohort from the COBRE dataset, comprising 72 patients with SZ and 75 HCs, was used for application. There were no significant differences in age or sex distributions between SZ and HC groups. The demographic statistics for the application cohort are summarized in **Supplementary Table S1**.

### IL-6-related components in the discovery cohort

Two ReHo components (ReHo IC4 and ReHo IC8) were significantly correlated with plasma IL-6 levels in the discovery cohort (ReHo IC4: *r* = 0.095, *p*_FDR_ = 0.049; ReHo IC8: *r* = 0.110, *p*_FDR_ = 0.027; Error! Reference source not found.**A**). No significant correlations were observed for components from the GMV modality. Next, the spatial maps of the two IL-6-related components were converted to Z-scores and displayed at |Z| > 3 (Error! Reference source not found.**B**). For ReHo IC4, positive associations were observed in the ventral visual stream (including the lingual gyrus and middle occipital gyrus) and the posterior cerebellum (Crus I/ Lobule VI). Negative weights were found in mPFC [the superior medial frontal gyrus and anterior cingulate cortex (ACC)], as well as the dorsal visual stream (cuneus) and bilateral supramarginal gyrus. ReHo IC8 exhibited positive associations in the default mode network (DMN) core hubs [bilateral precuneus, posterior cingulate cortex (PCC), mPFC], and angular gyrus) and the cognitive cerebellum (Crus I/II), alongside negative associations in the sensorimotor cortex (precentral and postcentral gyri), the dorsal attention network (dorsolateral superior frontal gyrus), and the motor cerebellum (Lobule VI). Detailed statistical results are summarized in **Supplementary Table S2**.

### Group differences of IL-6-component loadings between SZ and HC

To investigate the association between IL-6 brain patterns and brain abnormalities in SZ, we linearly projected the two IL-6-related ReHo components (ReHo IC4 and ReHo IC8) onto the SZ application dataset and computed subject-specific loading parameters. Two-sample *t*-tests were conducted to compare the component loadings between patients with SZ and HCs. Patients with SZ showed significantly higher loadings in ReHo IC8 (*t* = 3.73, *p* < 1 × 10□^3^) compared with HC (**Figure 2C**). In contrast, no significant group difference was found in ReHo IC4.

**Figure 1.**
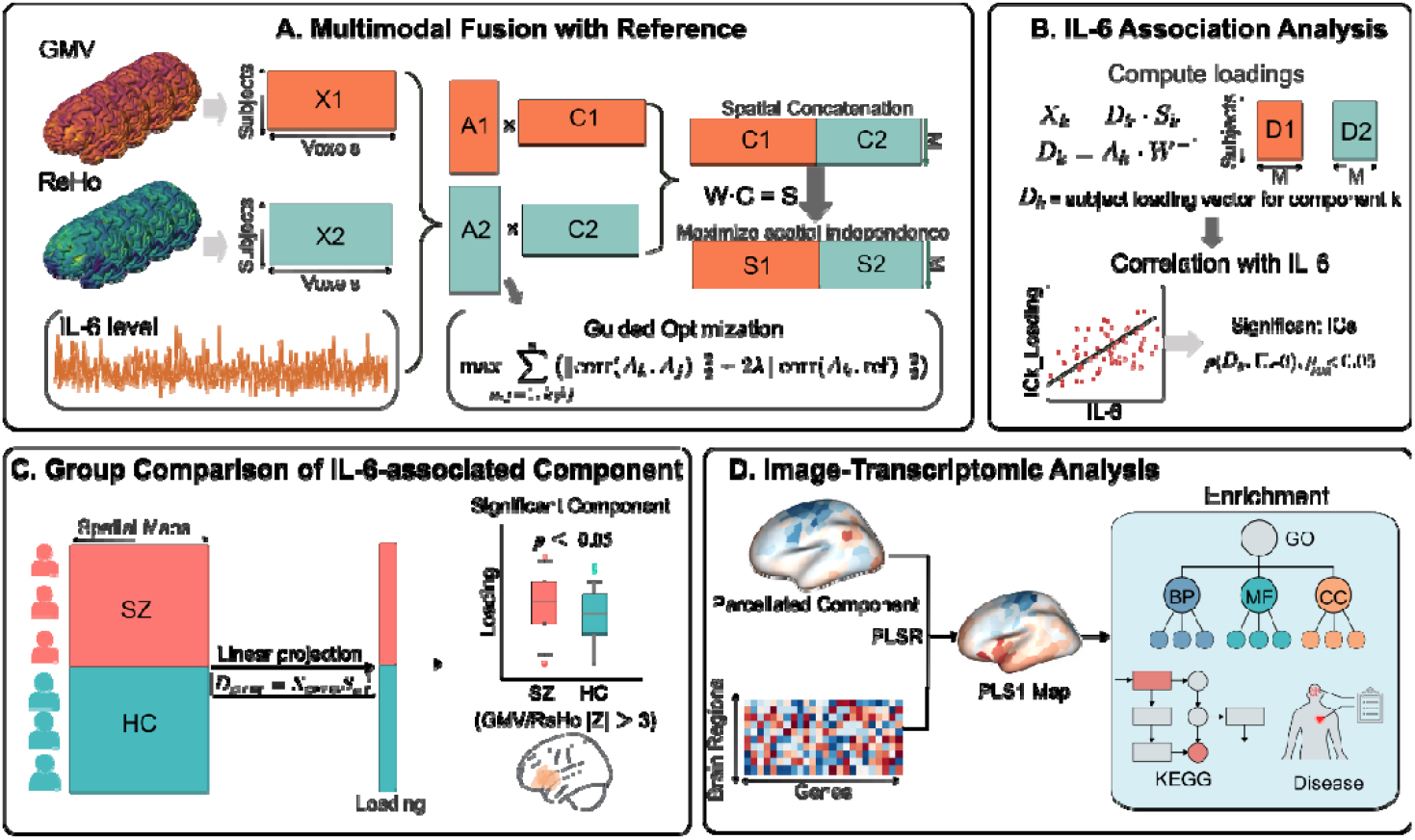
Schematic overview of the analysis framework. (A) Supervised Multimodal Fusion with Reference: GMV and ReHo maps from the discovery cohort were fused using MCCAR+jICA. Plasma IL-6 levels served as the reference signal to guide the optimization, maximizing the correlation between the mixing matrix (*A*_*k*_) and the reference vector while ensuring spatial independence of the components (*S*). **(B) IL-6 Association Analysis:** Subject-specific loading vectors (*D*_*k*_) were computed, and their associations with IL-6 levels were assessed. Components showing significant correlations (*p*_FDR_ < 0.05) were identified as inflammation-related ICs. **(C) Application in Independent Cohort:** The spatial maps of the identified IL-6-associated components were linearly projected onto an independent application cohort (SZ dataset). Group differences in component loadings between patients with SZ and HCs were tested to verify disease relevance. **(D) Image-Transcriptomic Analysis:** The spatial patterns of the validated components were aligned with cortical gene expression profiles from the AHBA using PLSR. Genes with significant spatial coupling (PLS1 weights) were subjected to functional enrichment analyses, including GO, KEGG pathways, and disease associations. Abbreviations: GMV, gray matter volume; ReHo, regional homogeneity; MCCAR+jICA, multi-site canonical correlation analysis with reference plus joint independent; IL-6, interleukin-6; *A*_*k*_, mixing matrix; *D*_*k*_, subject-specific loading vector; ICs, independent components; *p*_FDR_, false discovery rate–corrected p value; SZ, schizophrenia; HC, healthy controls; AHBA, Allen Human Brain Atlas; PLSR, partial least squares regression;PLS1, first latent component from PLS regression; GO, Gene Ontology; KEGG, Kyoto Encyclopedia of Genes and Genomes.

**Figure 2.**
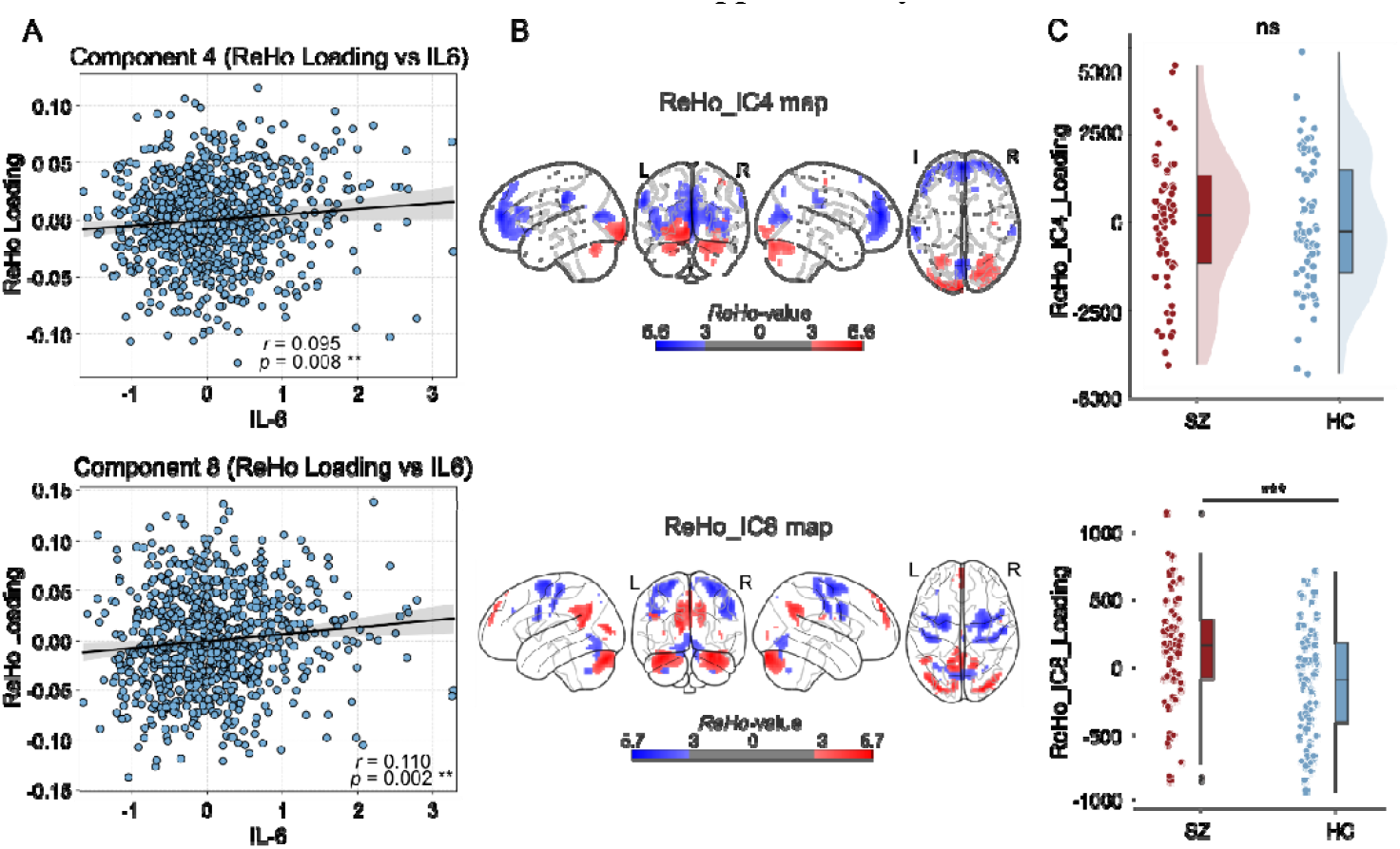
Identification and application of IL-6-associated components. (A) Associations with IL-6 in the Discovery Cohort: Scatter plots illustrating the significant linear correlations between plasma IL-6 levels and subject-specific ReHo loadings (y-axis) for Component 4 (top, *r* = 0.095, *p* = 0.008) and Component 8 (bottom, *r* = 0.110, *p* = 0.002) in the UK Biobank cohort. **(B) Spatial Distribution Maps:** The spatial patterns of ReHo IC4 and ReHo IC8, thresholded at |*Z*| > 3. Red (warm colors) indicates regions with positive weights (where ReHo increases with IL-6 levels), while blue (cold colors) indicates regions with negative weights (where ReHo decreases with IL-6 levels). **(C) Group Comparisons in the Application Cohort:** Comparison of component loadings between patients with SZ and HCs in the independent COBRE dataset. Abbreviations: IL-6, interleukin-6; ReHo, regional homogeneity; SZ, schizophrenia; HC, healthy controls; COBRE, Center for Biomedical Research Excellence dataset.

**Figure 3.**
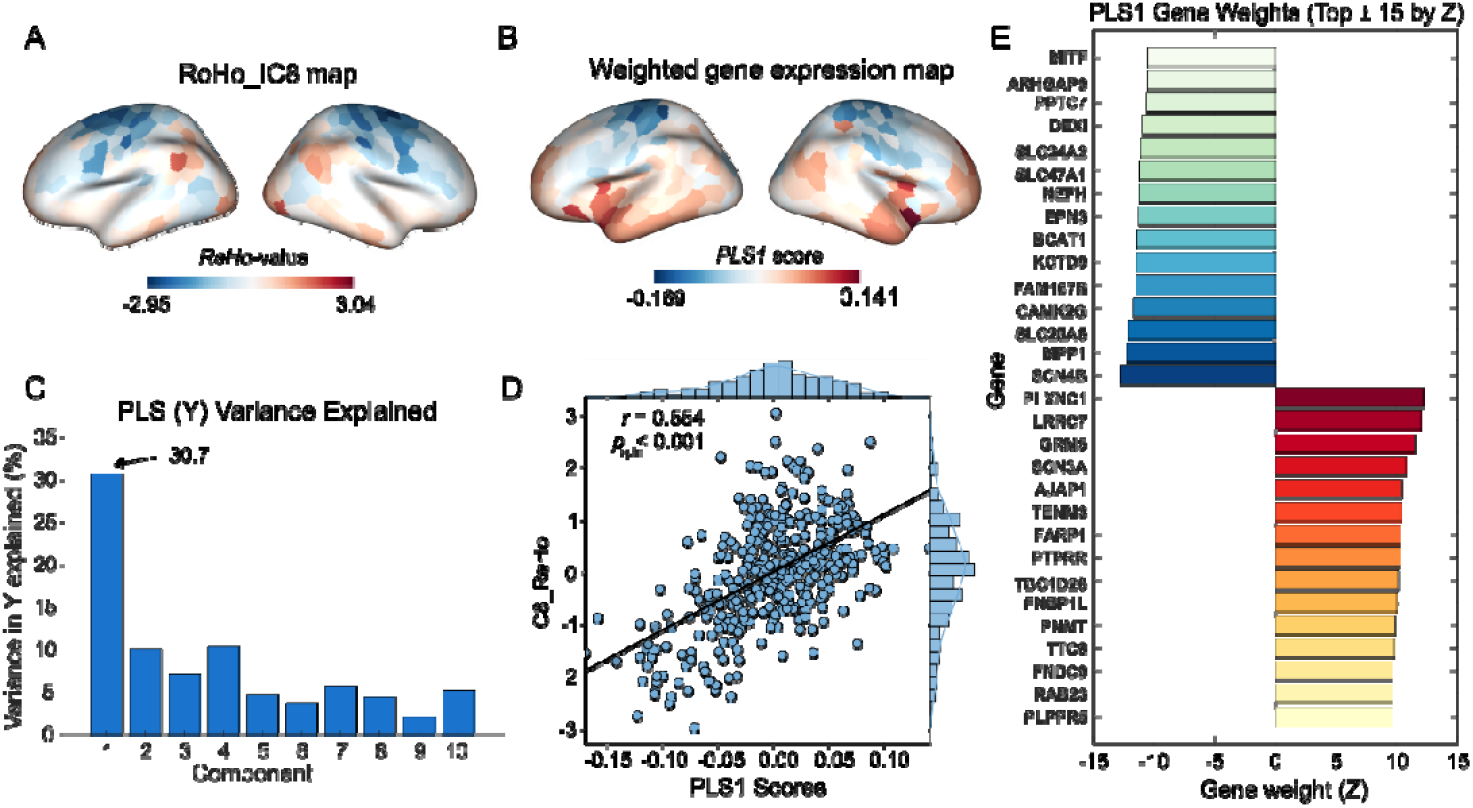
Spatial coupling between the IL-6-associated ReHo pattern and cortical gene expression profiles. **(A)** The spatial map of the disease-amplified ReHo component (ReHo IC8), resampled into the Schaefer-400 cortical parcellation. Red and blue indicate positive and negative loadings, respectively. **(B)** The spatial distribution of the first partial least Squares (PLS1) component scores derived from the Allen Human Brain Atlas (AHBA), representing the weighted gene expression pattern that best covaries with the ReHo IC8 map. **(C)** The percentage of variance in the ReHo IC8 map explained by each of the latent PLS components. The first component (PLS1) explained the largest proportion of variance (30.7%). **(D)** The scatter plot illustrating the significant spatial correlation between regional ReHo IC8 loadings (y-axis) and PLS1 gene scores (x-axis) across 400 cortical parcels. Statistical significance was determined using spatial permutation testing (spin test) to correct for spatial autocorrelation. **(E)** The top 15 genes with the highest positive (PLS1^+^, yellow-to-red) and negative (PLS1-, blue-to-green) weights contributing to the PLS1 component. Abbreviations: IL-6, interleukin-6; ReHo, regional homogeneity; PLS, partial least squares; PLS1, first latent component from partial least squares analysis; AHBA, Allen Human Brain Atlas;

**Figure 4.**
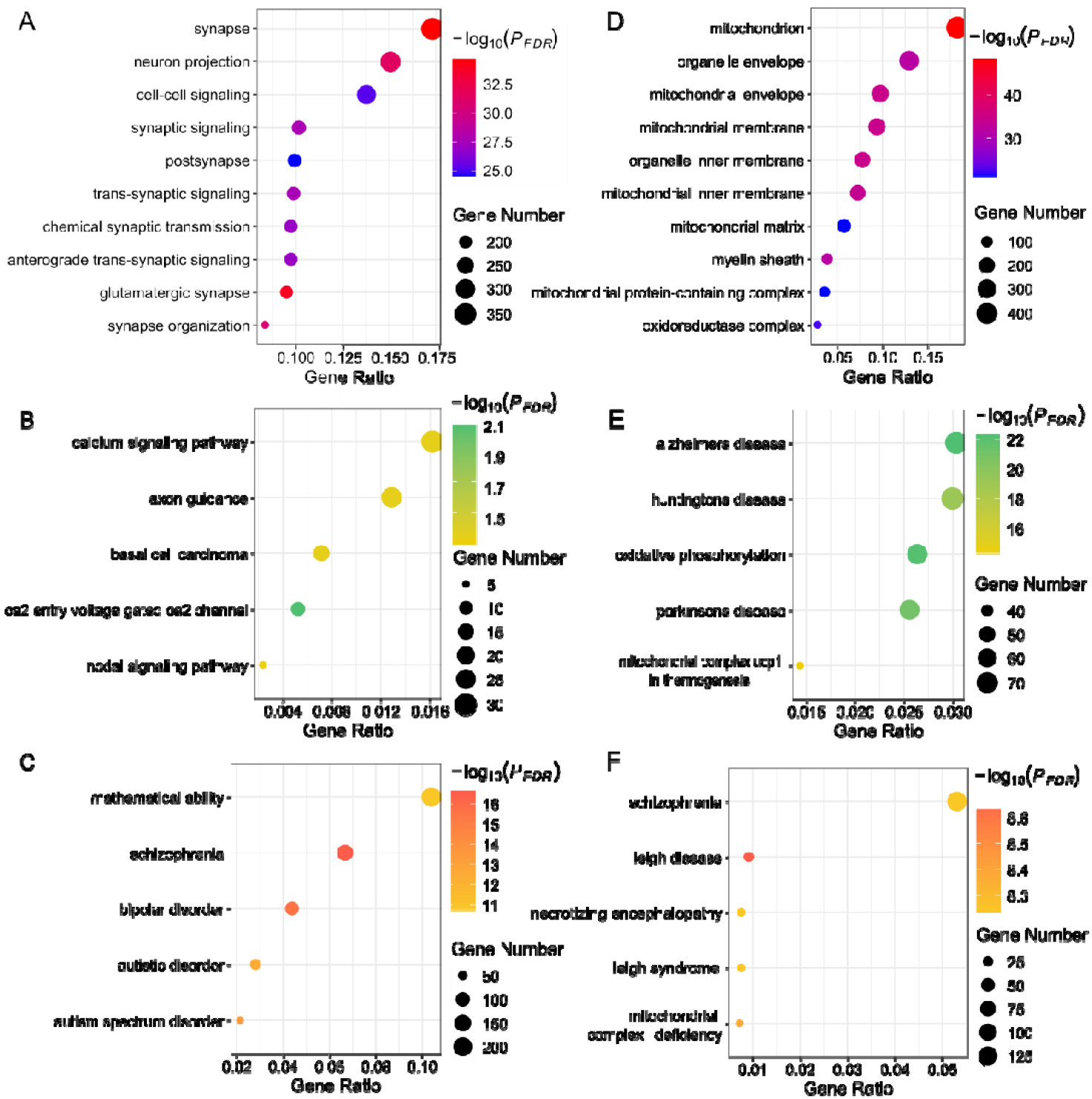
Functional enrichment analysis of the PLS1 gene signatures. (A, D) GO Enrichment: (A) Genes with positive weights were primarily enriched for synaptic processes, including synapse organization, neuron projection, and cell-cell signaling. (D) Genes with negative weights were significantly enriched for mitochondrial structures, such as the mitochondrial envelope and matrix. **(B, E) KEGG Pathway Enrichment:** (B) Positive weights were involved in neural signaling pathways, specifically calcium signaling and axon guidance. (E) Negative weights were enriched in oxidative phosphorylation and pathways related to neurodegenerative diseases (e.g., Alzheimer’s, Huntington’s). **(C, F) Disease and Trait Enrichment:** (C) Positive weights showed significant genetic overlap with cognitive traits (mathematical ability) and major psychiatric disorders (schizophrenia, bipolar disorder). (F) Negative weights were associated with mitochondrial disorders (Leigh disease/necrotizing encephalopathy) and schizophrenia. Abbreviations: PLS1, first latent component from partial least squares analysis; GO, Gene Ontology; KEGG, Kyoto Encyclopedia of Genes and Genomes; FDR, false discovery rate.

### Imaging-Transcriptome Association Analysis

To examine the spatial correspondence between the ReHo IC8 and cortical gene expression profiles derived from the Allen Human Brain Atlas, we performed PLS between ReHo and transcriptomic maps. The first latent component (PLS1) explained 30.7% of the variance in the imaging phenotype (**Error! Reference source not found**.). The spatial pattern of the PLS1 score map was significantly correlated with the ReHo IC8 score map (*r* = 0.55, *p*_*spin*_ < 0.001) (Error! Reference source not found.**D**). Genes were ranked according to their corrected PLS1 weights, and Z-scores were computed based on bootstrap-estimated standard errors. Genes with |*Z*| > 3 were included in the positive (PLS1^+^) and negative (PLS1^−^) gene lists, respectively (**Error! Reference source not found.E**).

To further reveal biological pathways involved in the PLS1^+^ and PLS1^−^ gene lists, we conducted GO and KEGG enrichment analyses. For the PLS1□ gene set, top GO terms were primarily related to synaptic organization and neuronal communication, such as synapse, neuron projection, cell–cell signaling, and chemical synaptic transmission (*p*_FDR_ < 0.05, **Error! Reference source not found.A**). Consistently, we found significant KEGG terms including neural signaling, including calcium signaling pathway and axon guidance (*p*_FDR_ < 0.05, **Error! Reference source not found.B**). Disease enrichment analysis indicated significant associations with major neuropsychiatric conditions (*p*_FDR_ < 0.05), including SZ, bipolar disorder, and autism spectrum disorder (**Error! Reference source not found.C**). For the PLS1□ gene set, enrichment shifted toward mitochondrial structure and bioenergetics: top GO terms included mitochondrion, mitochondrial membrane/envelope, organelle inner membrane, and mitochondrial matrix (**Error! Reference source not found.D**). KEGG pathways were related to mitochondrial energy metabolism and neurodegenerative terms (Parkinson’s, Huntington’s, and Alzheimer’s disease) (**Error! Reference source not found.E**). Disease enrichment further underscored mitochondrial dysfunction and clinical relevance, highlighting SZ alongside Leigh disease and related mitochondrial complex I/V deficiencies and infantile necrotizing encephalopathy (**Error! Reference source not found.F**). Given that ReHo IC4 appears to represent a conserved physiological response pattern with no significant group differences, its detailed imaging-transcriptomic results are provided in the Supplementary Materials (**Figure S2** and **Supplementary Results**).

## Discussion

In this study, we used the IL-6-guided supervised multimodal fusion approach to investigate the association between inflammation and brain network abnormalities in SZ. We found two distinct functional patterns associated with IL-6 levels. The first pattern (ReHo IC4), characterized by the mPFC and visual areas, appears to represent a conserved physiological response with no significant difference between the SZ and HCs. In contrast, the second pattern (ReHo IC8) was characterized by the default mode network and sensorimotor network, and represents a disease-amplified phenotype specific to SZ. Imaging-transcriptomic analyses further revealed that this pathological pattern was spatially coupled with gene expression profiles enriched for synaptic signaling and mitochondrial metabolism. Collectively, these results suggest that the IL-6 levels are related to the functional activities of specific brain networks, alterations of which were involved in the pathophysiology of SZ.

The ReHo IC4 weight map has a spatial pattern from the visual areas to the mPFC. The absence of a group difference between the SZ and HC cohort suggests that this neuro-immune coupling was preserved in patients, likely reflecting a fundamental physiological modulation of brain function by inflammation. This pattern suggests an inflammation-related shift toward externally driven sensory input and away from internally oriented prefrontal processing. Visual cortices primarily support the encoding of external environmental information, whereas the mPFC is central to self-referential and internal evaluative processes. The increased visual synchrony alongside mPFC suppression therefore likely reflects a physiological reweighting between external input and internal integration under inflammatory states.

The ReHo IC8 component was anchored in the sensorimotor areas and DMN, and showed positive deviations in SZ relative to HC. Hence, we inferred this component as a disease-relevant phenotype. This configuration reflects an inflammation-linked bias toward internally oriented mentation (DMN), accompanied by reduced synchrony in networks supporting externally directed sensory–motor processing. The DMN is centrally involved in internally oriented processes (Van Buuren et al., 2012), whereas sensorimotor networks are often implicated in deficits in external engagement and psychomotor function (Gao et al., 2019). The elevation of IC8 expression in SZ further indicates that this IL-6–associated functional configuration is amplified in the disorder. This pattern aligns with prevailing dysconnectivity models of SZ (Mlynek et al., 2025), which emphasize excessive dominance of internally oriented networks and reduced engagement of systems supporting interaction with the external environment. The elevated loading of ReHo IC8 in SZ suggests that peripheral inflammation preferentially engages large-scale networks already implicated in the disorder, accentuating an internal–external processing imbalance rather than producing a nonspecific global disruption.

In terms of the biological pathway of the ReHo IC8 component, the DMN-side of the pattern was associated with synaptic function, while the sensorimotor-side was associated with mitochondrial energy metabolism. Regions with positive loadings, i.e., the DMN spatially covaried with the expression of genes enriched for synaptic signaling and neuronal communication. This accords with preclinical evidence that IL-6 regulates synaptic plasticity and transmission (Gruol, 2015; Tancredi et al., 2000), suggesting that inflammation-related DMN hypersynchrony may be anchored in synaptic molecular architectures. Conversely, regions with suppressed ReHo loadings (sensorimotor cortex) were enriched for genes related to mitochondrial structure and oxidative phosphorylation. Given the growing evidence linking mitochondrial dysfunction to both inflammation and SZ (Morris & Berk, 2015; Rajasekaran et al., 2015), this spatial coupling suggests that the vulnerability of sensorimotor networks to inflammation-associated suppression might be related to local bioenergetic constraints. While these transcriptomic associations are correlational and derived from normative post-mortem atlases, they offer a plausible biological framework linking peripheral IL-6 to the macroscopic functional reorganization observed in ReHo IC8. Disease enrichment analyses further indicated that genes associated with this IL-6–linked functional pattern overlapped with major psychiatric disorders, including SZ, providing convergent support for the clinical relevance of the identified network.

Additionally, we found the distinct cerebellar regions involved in the two patterns. ReHo IC8 also involved cerebellar subregions in a functionally specific manner. In particular, the inflammation-linked pattern showed increased ReHo in cognitive Crus I but decreased ReHo in the motor Lobule VI, mirroring the cortical dissociation between default mode and sensorimotor networks. At a descriptive level, this intra-cerebellar dissociation is broadly consistent with the cognitive dysmetria account (Andreasen et al., 1999). For ReHo IC4, the involvement of posterior cerebellar regions may reflect a general inflammation-related coupling with perceptual systems, consistent with its conserved, non–disease-specific profile.

Several limitations should be considered. First, the study design is based on cross-sectional data and correlation analyses, which prevents causal inferences regarding inflammation and disease progression. Second, application in the SZ cohort assessed disease relevance by projecting the IL-6–associated pattern onto an independent dataset and testing group differences, but the absence of peripheral IL-6 measurements in this cohort prevented direct replication of inflammation–brain associations in patients. Next, the transcriptomic analysis relied on the AHBA dataset, which does not reflect the specific gene expression changes occurring in SZ patients. Thus, the imaging-transcriptomic analyses should be carefully considered, and future studies using patient-derived transcriptomic data are needed to validate these molecular associations. Finally, IL-6–associated effects were only found in functional local synchrony (ReHo), while not in GMV. These results may be attributed to the fact that peripheral inflammation influences brain structure in a more gradual and time-consuming manner than brain function. Further longitudinal studies using multiple structural features are needed to validate this inference.

## Conclusion

This study identified an IL-6–associated functional brain pattern that was selectively amplified in SZ, primarily reflected in intrinsic functional synchrony with a dissociation between default mode dominance and sensorimotor suppression. Imaging-transcriptomic analyses further revealed the molecular architecture underlying this pattern, coupling synaptic signaling with default mode regions and linking mitochondrial bioenergetic processes with the sensorimotor cortices. These findings provide a systems-level perspective on how peripheral immune signaling interacts with intrinsic brain networks in SZ.

## Code availability

The MCCAR+jICA algorithm has been implemented in the Fusion ICA Toolbox (FIT) and is available for download at https://trendscenter.org/software/fit.

## Acknowledgements

The authors declare that they have no conflict of interest. This work was supported by the National Natural Science Foundation of China (62403105, 62333003, 82121003, 62373079), the Sichuan Science and Technology Program (2026NSFSC1509), the China Postdoctoral Science Foundation (2023M740524), the Sichuan Province Innovative Talent Funding Project for Postdoctoral Fellows, the Medical Engineering Cooperation Funds from University of Electronic Science and Technology of China (ZYGX2021YGLH201), the Science and Technology Department of Sichuan Province (2025ZNSFSC0734), the Health Commission of Sichuan Province (24CXTD11), Health Commission of Chengdu (2024141), the Sichuan Preventive Medicine Association (SYYXHPT202420).

