## Supplement for "Inflammation in schizophrenia: Peripheral interleukin-6-related disease-specific functional activity abnormalities"

Running Title: Interleukin-6-related brain alterations in schizophrenia

Yun-Shuang Fan ^a,b#^, Jinxing Chen ^a,b#^, Liju Liu ^a,b^, Cuiling Zhang ^a,b^, Jing Guo ^a,b*^, Huafu Chen ^a,b*^, Mi Yang ^a*^

a The Clinical Hospital of Chengdu Brain Science Institute, School of Life Science and Technology, University of Electronic Science and Technology of China, Chengdu, China;

b MOE Key Lab for Neuroinformation, Brain-Computer Interface & Brain-Inspired Intelligence Key Laboratory of Sichuan Province, University of Electronic Science and Technology of China, Chengdu, China;

^#^ Co-first authors;

* Corresponding authors:

Mi Yang &

Huafu Chen, &

Jing Guo,

**Supplementary Table S1. Demographic characteristics of the participants in the application cohort (COBRE).**

| Characteristic | SZ (**n**=72) | HC (**n**=74) | *T* / χ^2^ Value | *P*-value |
| --- | --- | --- | --- | --- |
| Age (years) | 38.17 ± 13.89 | 35.82 ± 11.58 | t = -1.105 | 0.271 |
| Sex (M/F) | 58 / 14 | 51 / 23 | χ^2^ = 2.612 | 0.106 |

Note:

^a^Data are presented as mean ± standard deviation for continuous variables and counts for categorical variables.

^b^ Statistical comparison was performed using the independent two-sample t-test for age (equal variances not assumed due to Levene's test p < 0.05) and Pearson’s chi-square test for sex distribution.

Abbreviations: SZ, Schizophrenia; HC, Healthy Controls; M, Male; F, Female.

**Supplementary Table S2. Brain regions showing significant weights in the identified IL-6-associated components (ReHo_IC4 and ReHo_IC8).**

| Component | Anatomical Region (AAL3) | Hemisphere | Peak MNI (x, y, z) | Cluster Size (k) | Peak Intensity (Z) |
| --- | --- | --- | --- | --- | --- |
| ReHo_IC4 | **Ventral Visual Cortex** |  |  |  |  |
| (Positive) |  |  |  |  |  |
|  | Middle Occipital Gyrus / Lingual Gyrus | L | -6, -102, -9 | 249 | 5.15 |
|  | **Posterior Cerebellum** |  |  |  |  |
|  | Crus I / Lobule VI | R | 21, -84, -27 | 312 | 4.28 |
|  | Crus I / Lobule VI | L | -27, -66, -30 | 81 | 3.93 |
| (Negative) | **mPFC** |  |  |  |  |
|  | Superior Medial Frontal Gyrus / ACC | Bilateral | 0,54,15 | 1067 | -5.57 |
|  | **Dorsal Visual Cortex** |  |  |  |  |
|  | Cuneus / Superior Calcarine | L | 0, -69,12 | 166 | -4.67 |
|  | **IPL** |  |  |  |  |
|  | Supramarginal Gyrus | L | -60, -27,33 | 86 | -4.95 |
|  | Supramarginal Gyrus | R | 63, -27,30 | 30 | -3.6 |
| ReHo_IC8 | **Default Mode Network** |  | | | |
| (Positive) |  |  |  |  |  |
|  | Precuneus /PCC | L / R | -12, -54,18 | 253 / 88 | 4.96 |
|  | mPFC | L | 0,57,39 | 88 | 4.66 |
|  | Angular Gyrus | L | -45, -51,27 | 50 | 4.49 |
|  | **Posterior Cerebellum (Cognitive)** |  |  |  |  |
|  | Cerebellum Crus I / Crus II | R | 30, -90, -27 | 252 | 5.72 |
|  | Cerebellum Crus I / Crus II | L | -36, -84, -30 | 217 | 5.56 |
| (Negative) | **Sensorimotor / Dorsal Attention Network** |  |  |  |  |
|  | Superior Frontal Gyrus (Dorsolateral) / Precentral | R | 30, -6,63 | 422 | -5.15 |
|  | Superior Frontal Gyrus (Dorsolateral) / Precentral | L | -24, -6,69 | 198 | -5 |
|  | Postcentral Gyrus (Somatosensory) | L | -9, -24,60 | 135 | -4.58 |
|  | **Anterior Cerebellum (Motor)** |  |  |  |  |
|  | Cerebellum Lobule VI / Vermis | Bilateral | 0, -75, -6 | 199 | -4.06 |

Note:

^a^ Network assignment was determined based on the Yeo-7 network parcellation and anatomical location.

^b^ ReHo_IC4 and ReHo_IC8 denote the fourth and eighth independent components derived from the ReHo modality in the multimodal fusion analysis.

Abbreviations: MNI, Montreal Neurological Institute; AAL3, Automated Anatomical Labeling atlas version 3; mPFC, medial prefrontal cortex; ACC, anterior cingulate cortex; IPL, inferior parietal lobule; PCC, posterior cingulate cortex.

**Figure S1. Demographic characteristics of the discovery cohort from the UK Biobank.**
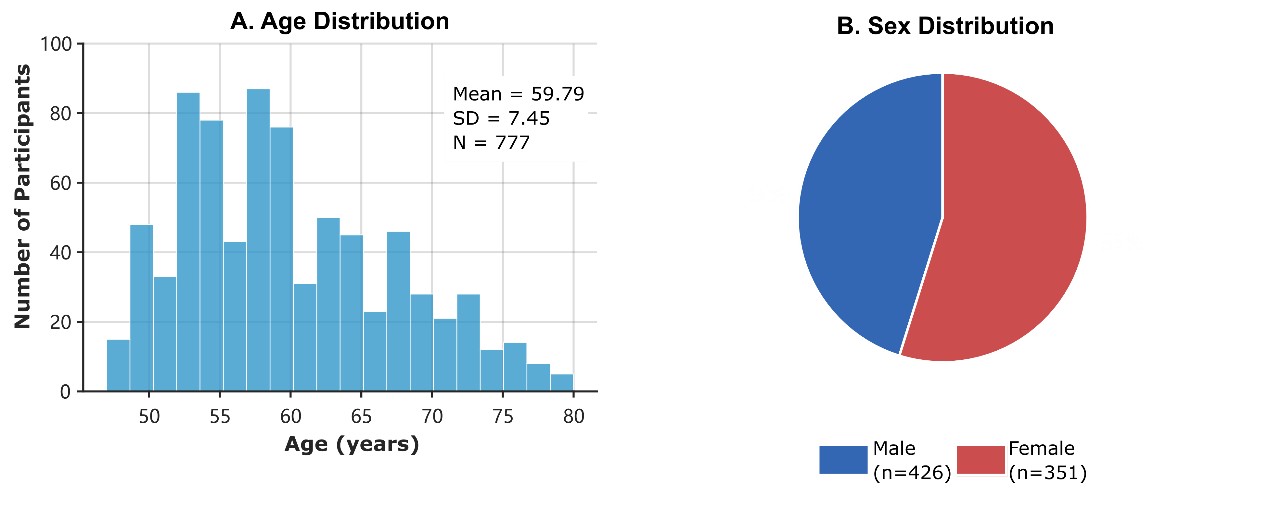

(A) Histogram illustrating the age distribution of the final sample included in the multimodal fusion analysis (*N* = 777). The study participants had a mean age of 59.79 ± 7.45 years.

(B) Pie chart representing the sex distribution of the cohort. The sample consisted of 426 males (55%) and 351 females (45%).

Note: The data represent the final participants after stringent quality control and exclusion criteria were applied.

**Figure S2. Functional enrichment analysis of the PLS1 gene signatures for ReHo_IC4.**

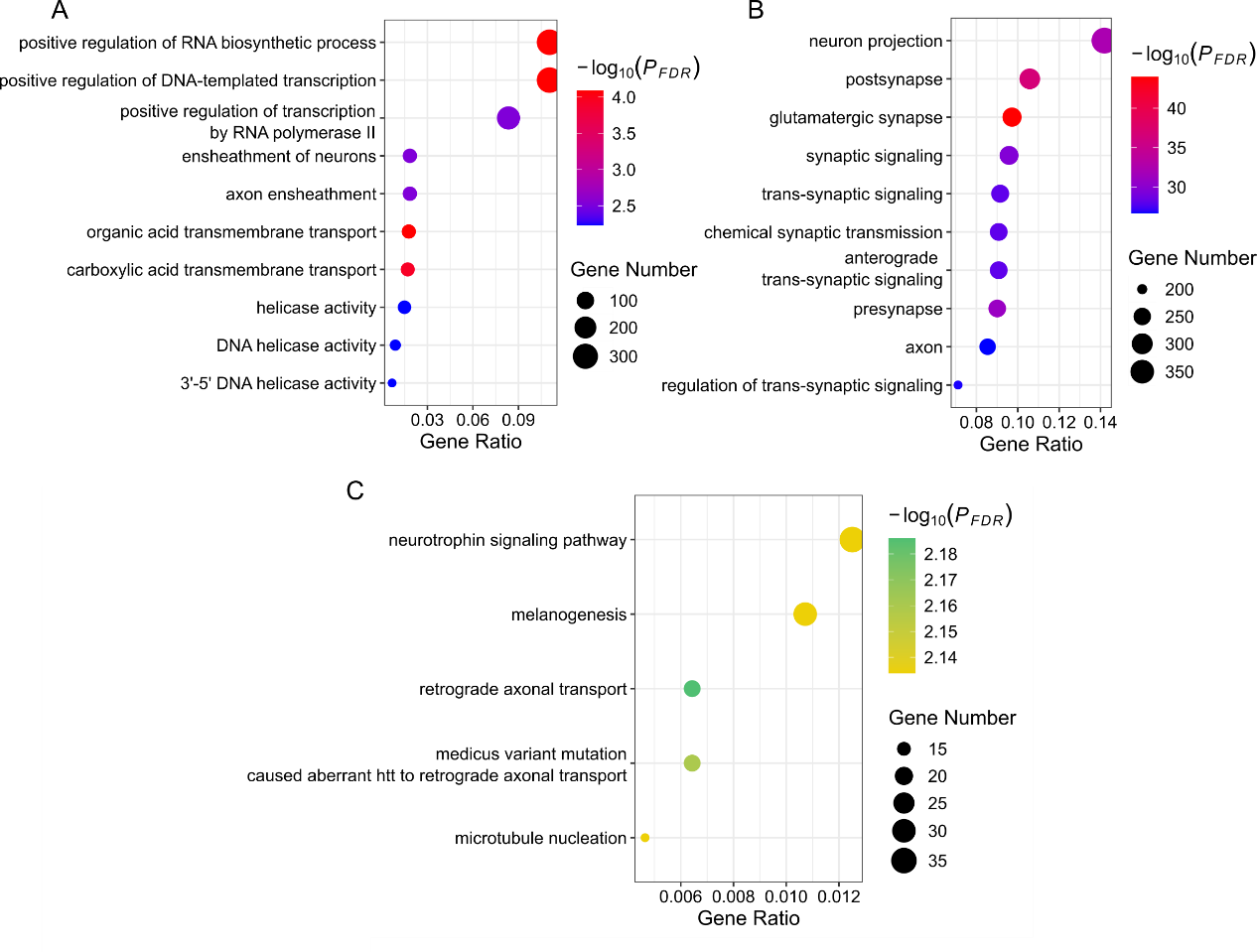

(A) GO enrichment analysis of genes with positive PLS weights (PLS^+^) revealed significant overrepresentation of transcriptional and biosynthetic processes, including positive regulation of RNA biosynthetic processes, DNA-templated transcription, and RNA polymerase II–mediated transcription, as well as axon and neuron ensheathment and transmembrane transport functions, indicating a role in fundamental cellular maintenance and structural support.

(B) GO enrichment analysis of genes with negative PLS weights (PLS^-^) demonstrated enrichment in neuronal and synaptic components and processes, such as neuron projection, pre- and postsynaptic structures, glutamatergic synapse, chemical and trans-synaptic signaling, and axonal regulation, highlighting close associations with synaptic communication and neural signaling.

(C) KEGG pathway enrichment analysis of PLS^-^ genes further identified pathways related to neurotrophic signaling, retrograde axonal transport, microtubule nucleation, and melanogenesis, suggesting involvement of cytoskeletal dynamics, axonal transport mechanisms, and neurotrophin-mediated signaling pathways underlying the PLS^-^ molecular profile.

Note: Dot size represents the gene count, and color indicates significance (*-log_10_*(*p*_FDR_)).

Abbreviations: PLS1, first latent component from partial least squares analysis; GO, Gene Ontology; KEGG, Kyoto Encyclopedia of Genes and Genomes; FDR, false discovery rate.

**Supplementary Results**

**Biological Characterization of the Physiological Component (ReHo_IC4)**

To reveal the molecular basis of ReHo_IC4, we also performed partial least squares (PLS) regression using the AHBA dataset. The first latent component (PLS1) explained 24.60% of the variance in the ReHo_IC4 spatial pattern. The spatial pattern of the PLS1 gene scores was significantly covaried with the ReHo_IC4 map (*r* = 0.50, *p*_spin_ < 0.001). To interpret the molecular basis of the conserved physiological pattern (ReHo_IC4), we performed gene enrichment analyses on the positive (PLS1^+^) and negative (PLS1^-^) gene weights derived from the regression (**Figure S2**).

Gene Ontology enrichment analysis showed that genes with positive PLS1 weights (PLS1^+^) were predominantly associated with fundamental cellular and transcriptional processes, including regulation of RNA biosynthesis, DNA-templated transcription, RNA polymerase II activity, and axonal ensheathment, suggesting a molecular profile related to cellular maintenance and structural support; notably, no KEGG pathways were significantly enriched for the gene set. In contrast, genes with negative PLS1 weights (PLS1^−^) were strongly enriched in neuronal and synaptic functions, encompassing neuron projection, pre- and postsynaptic structures, glutamatergic synapses, and trans-synaptic signaling. Consistently, Kyoto Encyclopedia of Genes and Genomes pathway analysis of PLS1− genes further highlighted neurotrophin signaling, axonal transport, and microtubule-related pathways, indicating that the ReHo_IC4 pattern is closely linked to molecular mechanisms underlying synaptic communication, axonal dynamics, and activity-dependent neural signaling.
